# Low abundant intestinal commensals modulate immune control of chronic myeloid leukemia stem cells

**DOI:** 10.1101/2024.05.31.595679

**Authors:** Magdalena Hinterbrandner, Francesca Ronchi, Viviana Rubino, Michaela Römmele, Tanja Chiorazzo, Catherine Mooser, Stephanie C. Ganal-Vonarburg, Kathy D. McCoy, Andrew J. Macpherson, Adrian F. Ochsenbein, Carsten Riether

**Affiliations:** Department of Medical Oncology, Inselspital, Bern University Hospital, University of Bern, Switzerland; Tumorimmunology, Department for BioMedical Research (DBMR), University of Bern, Bern, Switzerland; Graduate School of Cellular and Biomedical Sciences, University of Bern, Bern, Switzerland; Department of Visceral Surgery and Medicine, Inselspital, Bern University Hospital, University of Bern, Switzerland; Department for BioMedical Research, Visceral Surgery and Medicine, University of Bern, Switzerland; 6Charité - Universitaetsmedizin Berlin, corporate member of Freie Universität Berlin, Humboldt-Universität zu Berlin, and Berlin Institute of Health (BIH), Berlin, Germany: Institute of Microbiology, Infectious Diseases and Immunology (I-MIDI); Department of Physiology and Pharmacology, Snyder Institute of Chronic Diseases, Cumming School of Medicine, University of Calgary, Calgary, AB, Canada

**Author notes:** Corresponding author: Carsten Riether, Department of Medical Oncology, Inselspital, Bern University Hospital, University of Bern, Switzerland. M.H. and F.R. contributed equally to this paper.

**Keywords:** gut microbiota, immune escape, CML, LSC, Sutterella, anti-leukemic immunity

## Abstract

Leukemia stem cells (LSCs) are resistant to therapy and immune control. The reason for their resistance to elimination by cytotoxic T cells (CTLs) remains unclear. This study shows that specific low abundant Gram-negative intestinal commensals of the genus *Sutterella* suppress the anti-leukemia immune response in chronic myeloid leukemia (CML). We found that germ-free and specific opportunistic pathogen-free (SOPF) mice are protected from CML development and that colonization of SOPF mice with *Sutterella wadsworthensis*, but not other related and unrelated bacterial strains, rescues CML development. A higher prevalence of this microbe resulted in Myd88/TRIF-mediated CTL exhaustion in SPF compared to SOPF CML mice as evidenced by higher surface expression of exhaustion markers on CTLs, a reduced capacity to produce interferon-gamma and granzyme B and to kill LSCs *in vitro*. These findings provide new insights into the immune control of LSCs and identify *Sutterella* species as regulators of anti-leukemic immunity in CML.

## Introduction

Chronic myelogenous leukemia (CML) is associated with the Philadelphia (Ph′) chromosome, a reciprocal translocation between chromosomes 9 and 22, that results in formation of the oncogenic BCR-ABL1 fusion protein, a constitutively active tyrosine kinase that is necessary and sufficient for malignant transformation^1^. The BCR-ABL1 translocation occurs in hematopoietic stem or early progenitor cells known as leukemia stem cells (LSCs)^2^. The introduction of BCR-ABL1-targeting tyrosine kinase inhibitors (TKIs) has revolutionized the treatment of CML and greatly improved the prognosis of patients. However, only a subset of patients can successfully discontinue TKI therapy and maintain a treatment-free remission^3^. This is due to the persistence of TKI-resistant LSCs in the bone marrow (BM) of patients with CML^2^.

Clinical and experimental evidence suggests that CML induces leukemia-specific immunity that contributes to disease control^1^. Despite the fact that CML cells have a low mutational burden resulting in the generation of only a limited number of neo-antigens^4^, cytotoxic CD8^+^ T lymphocytes (CTLs) directed against leukemia antigens have been detected in the blood of CML patients^5^. Furthermore, CTLs have been shown to be able to eliminate leukemia cells and LSCs *in vitro*^6–9^. However, activated CTLs fail to eliminate LSCs *in vivo* and actually promote their expansion^8–11^.

The gut microbiota is a complex microbial community involved in a variety of beneficial host functions, including modulation of innate and adaptive immune responses of the gut-associated lymphoid tissue^12^. Consequently, any perturbation in the composition of the gut microbiota may have detrimental effects on human health, ultimately leading to various acute or chronic disease states^13^. Recent studies have reported that the gut microbiota promotes local inflammation and the development of gastrointestinal cancer^14^. In contrast, commensal microbes may exert opposing effects on tumours by priming the host immune system and enhancing the anti-tumour immune response^14^. Furthermore, the microbiota has an important role on the outcome of immunotherapeutic interventions in human cancer development^15–17^. Despite the accumulating evidence linking the microbiota to solid cancers, the role of the intestinal microbes in the initiation and development of hematological malignancies such as CML has not been thoroughly elucidated. In addition, only a few studies showed a role of the gut microbiota in the regulation of myelopoiesis during homeostasis and the development of immune myeloid cells^18–21^. However, ulcerative colitis patients with an altered microbiota have been reported to display an increased risk of developing myeloid leukemia^22^. It is not known whether this increased risk is caused directly by the leukemia or by the immunosuppressive treatment of colitis patients.

In the current study, we investigated host-microbiota interaction in CML development using a well-established murine retroviral transduction and transplantation model of BCR-ABL1-induced CML-like disease^6^. This model overall recapitulates the genetic and pathological features of human disease and enabled us to study the complex interaction between the intestinal commensal microbiota and the host immune system in the context of a developing leukemia. Using this model, we found that germ-free and SOPF mice are protected from CML development and that colonisation of SOPF mice with the Gram-negative bacterium *Sutterella wadsworthensis* (*S. wadsworthensis*) promotes CML development. A higher prevalence of this microbe in the intestine triggered CTL exhaustion in the bone marrow (BM) of specific-pathogen-free (SPF) CML mice. This resulted in a limited potential to eliminate LSCs and to protect from leukemia development. Colonization with S. wadsworthensis restored CML in SOPF mice. Mechanistically, we could show that Myd88/TRIF-mediated antigen recognition of intact bacterial cells, even when inactivated, on host immune cells but not circulating bacterial products or metabolites, is sufficient and necessary for the development of CML in vivo.

## Results

### The gut microbiota promotes CML development in SPF mice

We investigated the functional importance of the gut microbiota in the initiation and progression of CML in a well-established murine transplantation CML model^6^. Transplantation of 3x10^4^ BCR-ABL1-GFP-transduced lineage-negative (lin-)Sca-1^+^c-Kit^+^ BM cells (from here on termed LSC) into germ-free (GF) non-irradiated mice (GF CML) failed to induce CML and resulted in long-term survival of the mice while the disease regularly developed in mice kept under specific-pathogen free (SPF) conditions (SPF CML)**(Fig. 1A-C)**. No residual BCR-ABL1-GFP^+^ cells were detected by flow cytometry (FACS) in the blood, spleen and bone marrow (BM) of surviving GF CML mice 90 days after transplantation (data not shown). To determine residual disease using the most sensitive assay, we transplanted 5x10^6^ whole BM cells from surviving primary GF CML mice into lethally irradiated (2x6.5 Gy) SPF secondary recipients. All secondary recipients survived up to 90 days without evidence of leukemia **(Fig. 1D)**, suggesting that disease-initiating and -maintaining LSCs were eliminated or at least successfully controlled in these mice. The protection of GF mice from CML could not be explained by an impaired homing capacity of LSCs into the BM, as a similar number of functional LSCs were detected in the BM 14h after transplantation by FACS and colony-forming assays **(Fig. S1, A-C)**.

**Figure 1:**
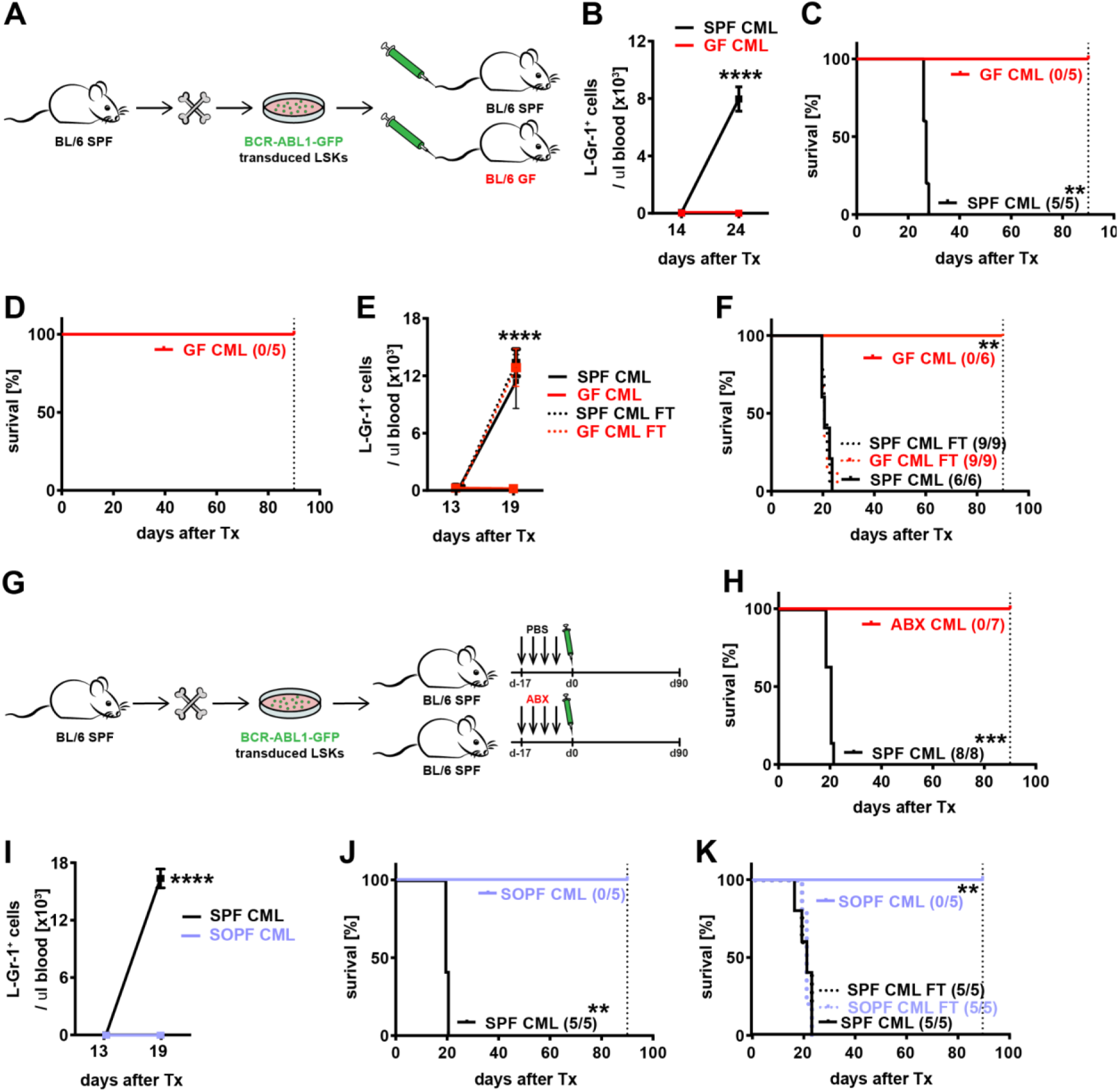
The gut microbiota contributes to CML development in mice. (A) Experimental plan. 3x10^4^ LSC were injected i.v. into nonirradiated C57BL/6 SPF (BL/6 SPF) and germ-free (GF) recipient mice and CML development was assessed. (B) Numbers of BCR-ABL1-GFP^+^ granulocytes/μl in blood (C) and Kaplan-Meier survival curves resulting from primary transplantations (Tx) of LSCs in C57BL/6 SPF and GF mice (n=5 mice/group). Representative data from at least two independent experiments are shown. Significance was determined using a two-way ANOVA followed by Bonferroni post-test (B) and a log-rank test (C). (D) Kaplan-Meier survival curve of secondary CML mice. BM cells of surviving primary CML mice were injected i.v. into lethally irradiated secondary BL/6 recipients, and survival was monitored (n=5 surviving GF CML mice). Representative data from at least two independent experiments are shown. Significance was determined using a log-rank test. (E) Numbers of BCR-ABL1-GFP^+^ granulocytes/μl in blood (F) and Kaplan-Meier survival curves resulting from primary transplantations (Tx) of 3x10^4^ LSCs in untreated C57BL/6 SPF and GF mice and SPF and GF mice undergoing previous fecal transfer (FT) from SPF mice (n=6-9 mice/group). Pooled data from two independent experiments are shown. Significance was determined using a two-way ANOVA followed by Bonferroni post-test (E) and a log-rank test (F). (G) Experimental plan. SPF mice were treated with PBS or antibiotics (ABX) for 17 days prior to CML induction and survival was monitored. (H) Kaplan-Meier survival curves resulting from primary transplantations (Tx) of 3x10^4^ LSCs in C57BL/6 SPF and ABX-treated mice (n=7-8 mice/group). Pooled data from two independent experiments are shown. Significance was determined using a log-rank test. (I) CML development and (J) survival of C57BL/6 SPF (SPF) and SOPF (SOPF) mice. Representative data from at least two independent experiments are shown. Significance was determined using a two-way ANOVA followed by Bonferroni post-test (I) and a log-rank test (J). (K) Kaplan-Meier survival curves resulting from primary transplantations (Tx) of LSCs in untreated C57BL/6 SPF and SOPF mice and SPF and SOPF mice undergoing previous fecal transfer from SPF mice (n=5 mice/group). One representative out of two independent experiments is shown. Significance was determined using a log-rank test. Data are represented as mean ± SD. **p < 0.01; ***p < 0.001; ****p < 0.0001.

To further confirm that the microbiota contributes to CML development in SPF colonized mice, we examined CML development in GF mice exposed to the SPF microbiota via co-housing (referred to as EX-GF mice) and fecal microbiota transfer (FT) prior to leukemia induction (referred to as FT-GF mice). EX-GF mice developed CML and succumbed to the disease with similar kinetics and latency as SPF CML mice **(Fig. S1D, E)**. Similarly, FT from SPF to GF mice completely restored CML development in recipient mice **(Fig. 1E, F)**.

Next, we depleted the gut microbiota by broad-spectrum antibiotic treatment and assessed the CML development. Similar to GF mice, SPF mice treated with broad-spectrum antibiotics^23^ were protected from CML development **(Fig. 1G, H)**.

To evaluate whether any type of stable microbial consortia with different complexities is sufficient to promote CML development, we transplanted LSCs into specific opportunistic pathogen-free (SOPF), and different gnotobiotic models such as sDMDMm (stable defined moderately diverse mouse microbiota)^24^ and Cuatro^25^ stably colonized mice. All gnotobiotic as well as SOPF mice were completely protected from CML development **(Fig. 1I, J and Fig. S2A, B)**. Similar to GF mice, FT from SPF mice was sufficient to restore susceptibility to CML in SOPF mice **(Fig. 1K)**.

Collectively, these results suggest that either a very complex and diverse microbiota or specific bacteria that are typical of the SPF microbiota but absent in the SOPF and gnotobiotic mouse models are required to initiate CML development.

### *Sutterella* and *Bilophila* strains are enriched in the feces of SPF mice and promote CML development

Since FT from SPF mice was sufficient to restore CML susceptibility in SOPF mice (**Fig. 1K**), we hypothesized that bacterial strains present in SPF mice and absent/low in SOPF mice may promote CML development. Therefore, we investigated the gut microbiota under the different hygiene conditions studied by 16S rRNA amplicon sequencing. SOPF and SPF mouse strains were from the same genetic background, obtained from the same supplier and fed the same diet. The analysis revealed that the gut of SPF mice harbored distinct commensal bacterial communities compared to the gut of SOPF mice **(Fig. 2A-C, Table S1-4)**. Alpha-diversity was very similar between the different hygiene conditions (**Fig. S3A, B**). The most common bacterial strains present at comparable levels in the gut of naive SPF and SOPF mice included *Bacteroidetes, Firmicutes* and *Actinobacteria* **(Fig. 2B-C)**. Bacteria from the phyla *Proteobacteria, Deferribacteres* and *Verrucomicrobia* were significantly overrepresented, whereas *Tenericutes* phylum was underrepresented in SPF compared to SOPF mice **(Fig. 2B-C)**. Among the *Proteobacteria* phylum, several bacterial genera were significantly overrepresented in SPF mice **(Fig. 2D, Tables S1-4)**. Among these, some consist in the gram-negative, anaerobic and non-spore-forming bacteria of the genus *Sutterella*^26^ and the gram-negative, obligately anaerobic and bile-resistant bacteria of the genus *Bilophila*^27^. Similar results in terms of alpha- and beta-diversity have been obtained also via metagenomics analysis **(Fig. S3C-E)**. The metagenomics analysis also revealed that no other archaeal or micro-eukaryotic species were found present in our samples **(Table S5)**.

**Figure 2:**
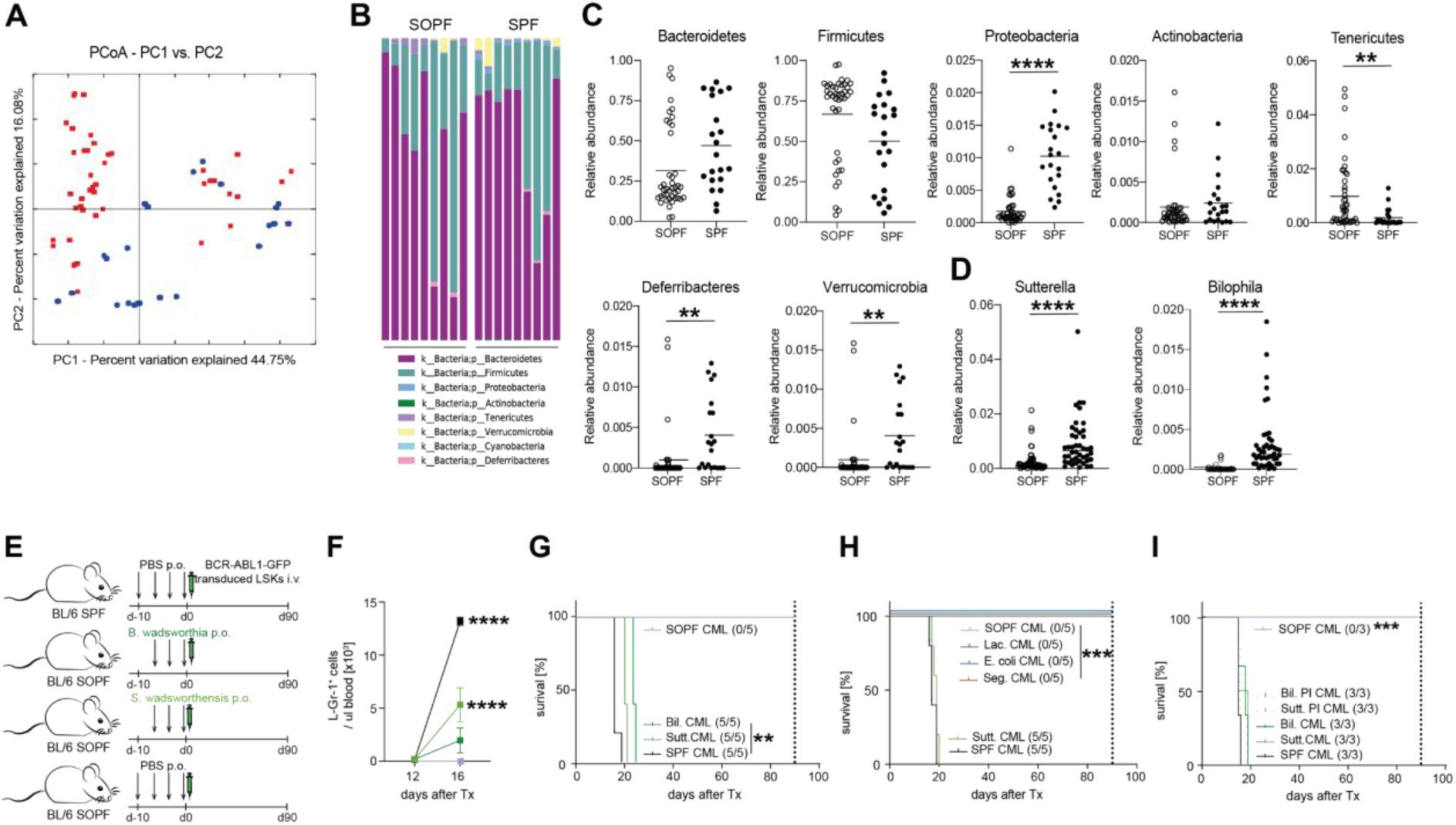
*Sutterella* and *Bilophila* strains promote CML development. (A-B) Microbiota composition analysis with 16S rRNA amplicon sequenicng with ionTorrent. (A) Principal coordinate analysis (PCoA). The PCoA PC1 and PC2 dimensions represent 60.83% of the microbiome variation between SPF and SOPF mice. Weighted UniFrac beta-diversity distance analysis between C57BL/6 SOPF and SPF fecal microbiota (ANOSIM statistical comparison) was significant, p-value 0.0001, number of permutations 9999. (B-D) Relative abundances of indicated bacteria taxa in feces from C57BL/6 SOPF and SPF mice (n=44 and 23, respectively). (B) Stool microbial composition at the phyla level in adult C57BL/6 SOPF and SPF mice. Each bar represents a single mouse. (C) Relative abundances of indicated bacterial phyla in feces from C57BL/6 SOPF and SPF mice. Each dot represents a single mouse. (D) Relative abundances of indicated bacterial genera in feces from C57BL/6 SOPF and SPF mice. Each dot represents a single mouse. (E) Experimental plan. C57BL/6 SOPF mice were colonized with *S. wadsworthensis* and *B. wadsworthia* at days -10, -7, -3, and 0 prior to CML induction. (F) Numbers of BCR-ABL1-GFP^+^ granulocytes/μl in blood (E) and Kaplan-Meier survival curves (G) resulting from primary transplantations (Tx) of 3x10^4^ LSCs in mice previously mono-colonized with *S. wadsworthensis* (Sutt. CML) and *B. wadsworthia* (Bil. CML) C57BL/6 SOPF mice and control C57BL/6 SOPF and SPF mice (n=5 mice/group). Representative data from at least two independent experiments are shown. Significance was determined using a two-way ANOVA followed by Bonferroni post-test (F) and a log-rank test (G). (H) Kaplan-Meier survival curves of SOPF mice colonized with *S. wadsworthensis* (Sutt. CML), *L. reuteri* (Lac. CML), S.copri (Seg. CML) and *E. coli* (E. coli CML) (n=5 mice/group). Representative data from at least two independent experiments are shown. Significance was determined using a log-rank test. (I) Kaplan-Meier survival curves of SOPF CML mice colonized with live and peracetic acid-inactivated bacteria (PI) *S. wadsworthensis* and *B. wadsworthia* (n=3 mice/group). Representative data from at least two independent experiments are shown. Significance was determined using a log-rank test. Data are represented as mean. **p < 0.01; ***p < 0.001; ****p < 0.0001.

To further evaluate whether these bacterial strains were able to trigger CML development, SOPF mice were colonized with live *Sutterella wadsworthensis (S. wadsworthensis)* and *Bilophila wadsworthia (B. wadsworthia)*. Colonization of SOPF mice with *S. wadsworthensis* and *B. wadsworthia* increased the relative abundance of these bacterial species to the level observed in SPF mice (**Fig. S4A)**. In addition, oral gavages with either bacterial strain rendered SOPF susceptible to CML development **(Fig. 2E-G)**. These differences in CML development could not be attributed to different steady-state levels or changes in bacterial biomass after administration of *S. wadsworthensis* (**Fig. S4B)**. Interestingly, colonization with *S. wadsworthensis* and *B. wadsworthia* did not induce CML in GF and sDMDMm mice **(Fig. S5A, B)**.

To determine whether colonization with phylogenetically *Sutterella-*related and unrelated bacterial strains was sufficient to promote CML development independently of the presence of *S. wadsworthensis* and *B. wadsworthia*, we transplanted LSCs into SOPF mice supplemented with different bacteria. To understand if the supplementation of other members of the *Proteobacteria* phylum would be sufficient to trigger CML development, we colonized SOPF mice with the *E. coli HS*^28^. To test if the complementation of any bacterium that was underrepresented in SOPF compared to SPF would trigger CML development, we gavaged the gram-negative *Bacteroidetes* strain *Segatella copri* (*S. copri*, DSM 18205) or the gram-positive, aerobic *Firmicutes Limosolaactobacillus reuteri* (*L. reuteri* I49^29^) in SOPF mice. *Prevotellaceae* and *Lactobacillales* were indeed significantly decreased in abundance in SOPF compared to SPF stools **(Table S1)**. Despite the fact that colonization with *L. reuteri*, *S.copri* and *E. coli HS* increased their relative abundance in SOPF mice, leukemia development was not affected by their colonization **(Fig. 2H, Fig. S4A, C)**.

### *S. wadsworthensis* bacterial antigens are sufficient and necessary for the development of CML in SPF mice

To investigate the mechanism through which *S. wadsworthensis* promotes CML development, we provided the bacterium in different inactivated forms. Firstly, the bacteria were inactivated by peracetic acid treatment prior to oral gavage to SOPF mice. Peracetic acid has previously been shown to induce high-titer specific intestinal IgA in the absence of measurable inflammation or species invasion^30^. Peracetic acid-inactivated (PI) bacteria were still able to promote CML development **(Fig. 2I)**. In contrast, gavage of heat-killed (HK) *S. wadsworthensis* did not contribute to disease development in SOPF mice **(Fig. S6A)**. By oral gavage of fluorescein isothyocyanate (FITC)-labeled dextran into SOPF and SPF mice, we could exclude altered systemic penetration of microbial products or microbes in steady state conditions **(Fig. S6B)**. Consistent with these findings, repeated intravenous injection (i.v.) of sera from SPF mice did not restore CML development in SOPF mice **(Fig. S6C)**. Collectively, these results suggest that specific intact bacterial cells, even if inactivated, but not circulating bacterial products or metabolites, are necessary for the development of CML *in vivo*.

### Colonization with *B. wadsworthia* increases the abundance of *Sutterella* bacteria in the gut

Bacteria–bacteria interactions within the microbiota have been shown to determine their growth and prevalence^12^. To understand why only colonization with *S. wadsworthensis* and *B. wadsworthia* affects CML development in SOPF mice, we determined how colonization with one bacterium affects the prevalence of the other bacteria in the gut and vice versa. 16S rRNA sequencing revealed that colonization with *B. wadsworthia* induced the accumulation of *Sutterella* bacteria in SOPF mice. In contrast, colonization with *S. wadsworthensis* did not affect the prevalence of *Bilophila* bacteria in SOPF mice **(Fig. S4A)**. These results suggest that the observed effect on CML development is most likely mediated by colonisation with *S. wadsworthensis*. As a result, we focused on the role of *S. wadsworthensis* in all future experiments.

### Myd88/Trif signaling in host cells mediates CML development

To understand how bacteria might promote CML development, we investigated whether microbial sensing through Toll-like receptors (TLRs) is involved. TLRs are sensors of bacterial-derived signals and mediate their downstream effects via the adaptor molecules MyD88 and TRIF^31^. To determine the contribution of host cell TLR-sensing to CML development, MyD88/TRIF-competent LSCs were transferred into *Myd88*^+/+^*/TRIF*^+/+^ (WT) or *MyD88^-/-^/TRIF^-/-^* double-knockout (KO) SPF littermate recipient mice (WT > WT and WT > KO CML). Similar to GF and SOPF mice, SPF *MyD88^-/-^/TRIF^-/-^*^-^ mice did not develop CML **(Fig. 3A-C)**. In complementary experiments, we transplanted *MyD88/TRIF*-double KO LSCs into *Myd88*^+/+^*/TRIF*^+/+^ and *MyD88^-/-^/TRIF^-/-^* SPF littermate recipient mice (KO > WT and KO > KO CML). Similar to *Myd88*^+/+^*/TRIF*^+/+^ CML cells, *MyD88^-/-^/TRIF^-/-^* CML cells only induced CML when transplanted into *Myd88/TRIF*-proficient recipients **(Fig. 3D)**, suggesting that sensing of bacterial signals by host cells is critical for CML development *in vivo*.

**Figure 3:**
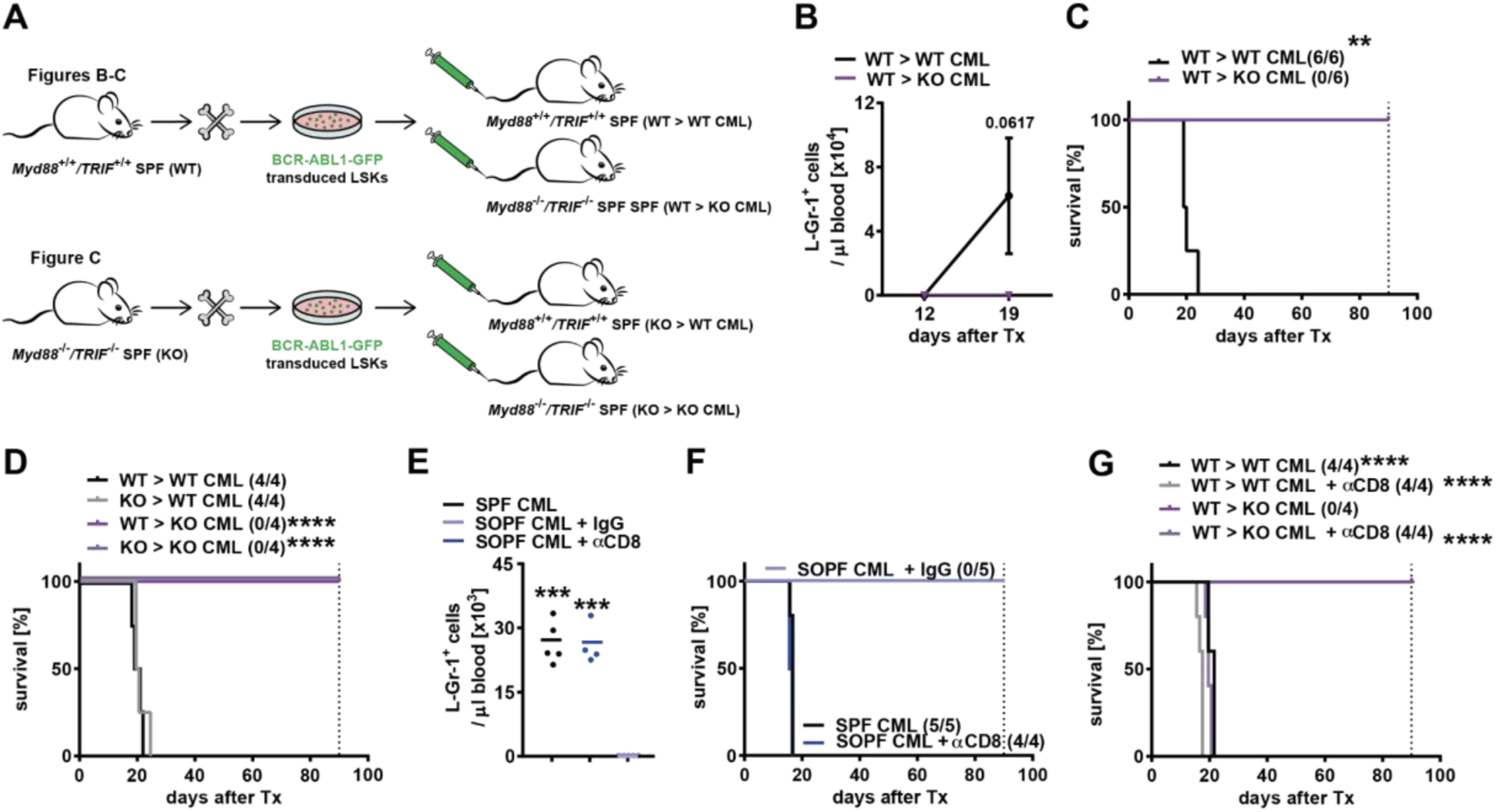
The intestinal microbiota contributes to the CML development through modulation of Myd88/TRIF signaling and CTLs. (A-D) Experimental plan. 3x10^4^ *MyD88/TRIF*-competent (WT) and - deficient (KO) LSCs were injected in *MyD88^+/+^/TRIF^-+/+^* and MyD88^-/-^/TRIF^-/-^ (WT > WT WT > KO, KO > WT and KO > KO CML, respectively) and CML development was assessed. (B-C) Numbers of BCR-ABL1-GFP**^+^** granulocytes/μl in blood (B) and Kaplan-Meier survival curves (C) resulting from primary transplantations (Tx) of 3x10^4^ WT LSCs into *MyD88/TRIF* WT and double-KO mice (n=6 mice/group). Representative data from at least two independent experiments are shown. Significance was determined using a two-way ANOVA followed by Bonferroni post-test (B) and a log-rank test (C). (D) Kaplan-Meier survival curves resulting from primary transplantations (Tx) of 3x10^4^ *MyD88/TRIF* WT and KO LSCs into *MyD88/TRIF* WT and KO mice (n=4 mice/group). Representative data from at least two independent experiments are shown. Significance was determined using a log-rank test. (E-F) Numbers of BCR-ABL1-GFP^+^ granulocytes/μl in blood (E) Kaplan-Meier survival curves (F) resulting from primary transplantations (Tx) of BL/6 LSCs into IgG and aCD8-treated SOPF mice and SPF control mice (n=4-5 mice/group). Representative data from at least two independent experiments are shown. Significance was determined using a log-rank test. (G) Kaplan-Meier survival curves resulting from primary transplantations (Tx) of 3x10^4^ *MyD88/TRIF*-competent LSCs into IgG and αCD8-treated *MyD88/TRIF*-competent (WT) and -deficient (KO) (n=4 mice/group). Representative data from at least two independent experiments are shown. Significance was determined using a log-rank test. Results are illustrated as mean. **p < 0.01; ***p < 0.001; ****p < 0.0001

### Cytotoxic α,β-CD8^+^ T cells protect SOPF mice from CML development

Different bacterial strains have been shown to regulate CD8^+^α,β CTL-mediated anti-tumor immunity in solid tumor models^12^. In addition, CTLs have been identified as key effector T cells in the control of LSCs in CML^5,6^. Therefore, we next determined whether CTLs contribute to the resistance of SOPF mice to CML development. Depletion of CTLs by monoclonal antibody treatment rendered the SOPF mice susceptible to CML development **(Fig. 3E, F)**. Similarly, GF mice treated with a depleting αCD8 antibody or GF lacking T and B cells (*Rag1*^-/-^ mice) developed the disease with a latency comparable to their SPF counterparts **(Fig. S7A, B)**.

Similarly, CD8^+^ T cell depletion restored CML development in *MyD88^-/-^/TRIF^-/-^* mice without affecting CML development in littermate controls **(Fig. 3G)**. Collectively, these data suggest that the gut microbiota contributes to CML development through modulation of Myd88/TRIF signaling and CTLs.

### *S. wadsworthensis* promotes activation of CTLs in the Peyers patches and in the BM under steady state conditions

*Sutterella* spp. are abundant in the duodenum with a decreasing gradient towards the colon^31^. Therefore, we hypothesized that *S. wadsworthensis* may affect the number or activation state of CTLs in the small intestine or secondary lymphoid structures in the gut during homeostasis. We therefore analyzed CTLs in Peyer’s patches (PP), mesenteric lymph nodes (MLN) and spleen in naive SPF and SOPF mice by flow cytometry **(Fig. 4A-C, fig. S6)**. The number of CTLs did not differ significantly between naive SPF and SOPF mice in any of the organs analyzed **(Fig. 4A)**. However, the number of activated CD44^+^ CTLs was significantly increased in the PP, but not in the MLN and spleen of SPF mice, compared to SOPF mice **(Fig. 4B)**. Similarly, TNFα single-producing and IFNℽ/TNFα double-producing CD44^+^ CTLs were significantly increased in frequency and absolute numbers in the PP, but not MLN and spleen of SOPF mice **(Fig. 4C-G)**. Furthermore, we found a 5-fold increase in CX3CR1^+^ CD44^+^ CTLs expressing programmed death 1 (PD-1) in the PP of SPF mice compared to SOPF mice **(Fig. 4H, I)**. Finally, we investigated whether colonization with *S. wadsworthensis* in SOPF mice affected cytokine secretion of CTLs in the PP. Intracellular FACS staining revealed that coloniation with *S. wadsworthensis* increased the number of TNFα single and IFNℽ/TNFα double producing CD44^+^ CTLs by a factor of 3 and 2.5, respectively **(Fig. 4J, K).**

**Figure 4:**
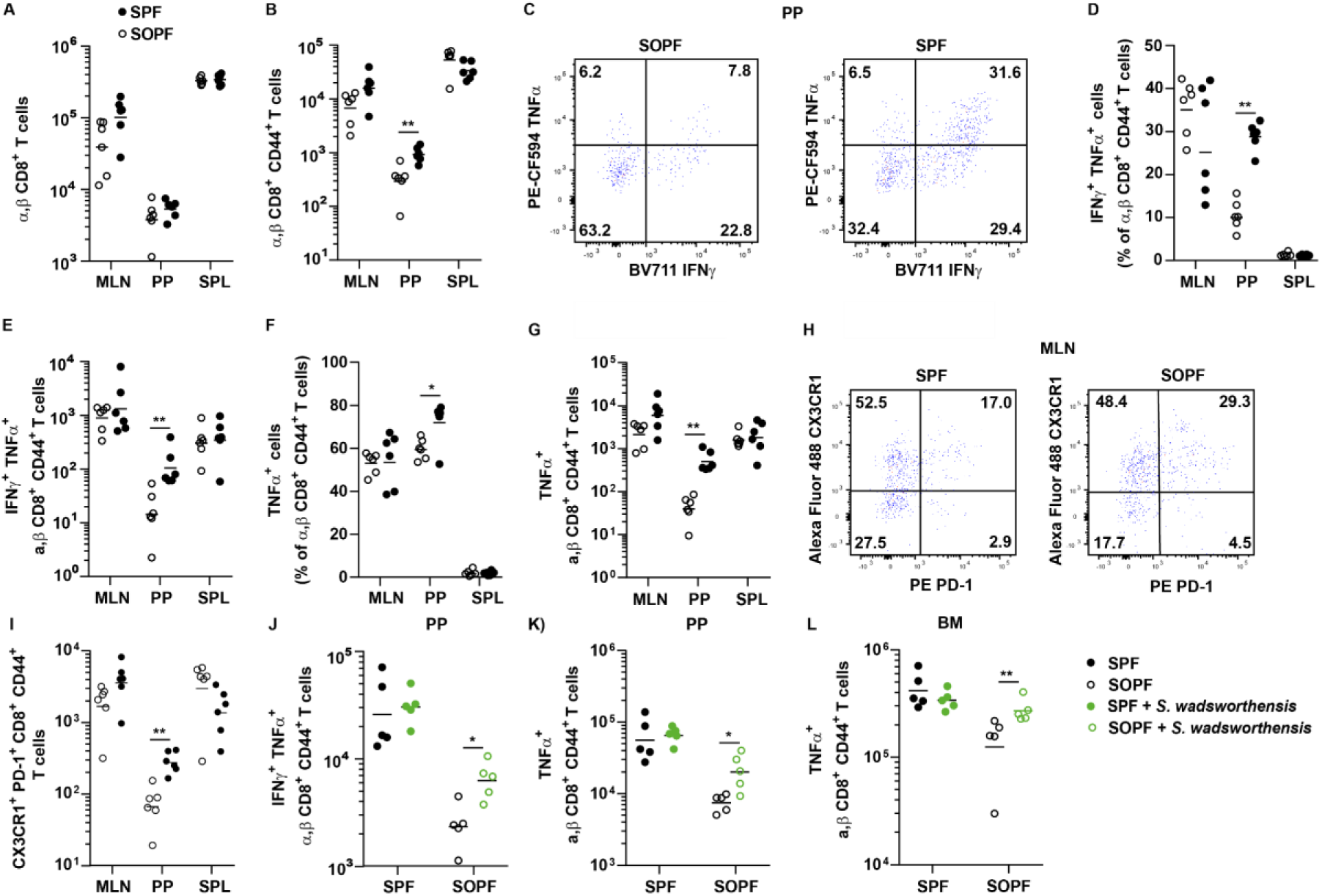
*S. wadsworthensis* induces IFNℽ and TNFα secretion of CTLs in Peyers patches. Numbers of (A) CTLs, (B) CD44^+^ CTLs in mesenteric lymph nodes (MLN), Peyers Patches (PP) and spleen of SPF and SOPF mice (n=6 mice/group). Representative data from at least two independent experiments are shown. Significance was determined using a Mann Whitney test. (C) Representative FACS plot for intracellular FACS stainings for IFNℽ and TNFα production by CTLs in SPF and SOPF mice. (D) Frequencies and (E) absolute numbers of IFNℽ and TNFα-double producing CTLs in MLN, PP and spleens of SPF and SOPF mice (n=6 mice/group). Representative data from at least two independent experiments are shown. Significance was determined using a Mann Whitney test. (F) Frequencies and (G) absolute numbers of TNFα producing CTLs in MLN, PP and spleens of SPF and SOPF mice (n=6 mice/group). Representative data from at least two independent experiments are shown. Significance was determined using a Mann Whitney test. (H) Representative FACS plot for PD-1 and CX3CR1 expression on CTLsin SPF and SOPF mice. (I) Absolute numbers of CX3CR1**^+^**PD1**^+^** CTLs in MLN, PP and spleens of SPF and SOPF mice (n=6 mice/group). Representative data from at least two independent experiments are shown. Significance was determined using a Mann Whitney test. (J-L) Absolute numbers of (J) IFNℽ and TNFα-double and (K) TNFα-producing CTLs in PP and TNFα-producing CTLs in BM of naive SPF and SOPF mice and SPF and SOPF colonized with *S. wadsworthensis* (n=5 mice/group). Representative data from at least two independent experiments are shown. Significance was determined using a Mann Whitney test. Results are shown as mean. *p < 0.05; **p < 0.01.

In CML, LSCs are mainly located in the BM and the spleen^2,32^. Therefore, we next determined the number of TNFα-producing CTLs in the BM and spleen in SPF and SOPF mice. TNFα-producing CTLs were increased in the BM but not in the spleen of SPF mice **(Fig. 4K and data not shown)**. Colonization of SOPF mice with *S. wadsworthensis* restored the number of TNFα-producing BM CD44^+^ CTLs to comparable levels of SPF mice. Overall, these data suggest that colonization with *S. wadsworthensis* leads to the activation of CTLs in PP and the BM.

### The gut microbiome modulates anti-leukemic immune CTL responses in CML

We next investigated whether and how the gut microbiome affects anti-leukemic immune CTL responses. We first evaluated LSC numbers in SPF and GF mice at different time points after leukemia induction. LSC numbers were comparable in GF and SPF mice 3 days after CML induction. In contrast, LSC numbers were significantly reduced in the BM of GF mice 6 days after induction. The reduction of LSCs in GF mice was dependent on the presence of CTLs (**Fig. 5A**). Similar to GF mice, LSKs in the BM of SOPF mice were reduced 6 and 9 days after transfer of CTV-labeled LSKs compared to the BM of SPF mice (**Fig. 5B**). The reduction in LSKs could not be attributed to changes in the cell cycle, as indicated by comparable frequencies of CTV-positive LSCs and a similar amount of cell divisions 6 and 9 days after injection (**Fig. 5C, D** **and data not shown)**. These findings suggest that the microbiome contributes to the control of CML by modulating CTLs in the BM.

**Figure 5:**
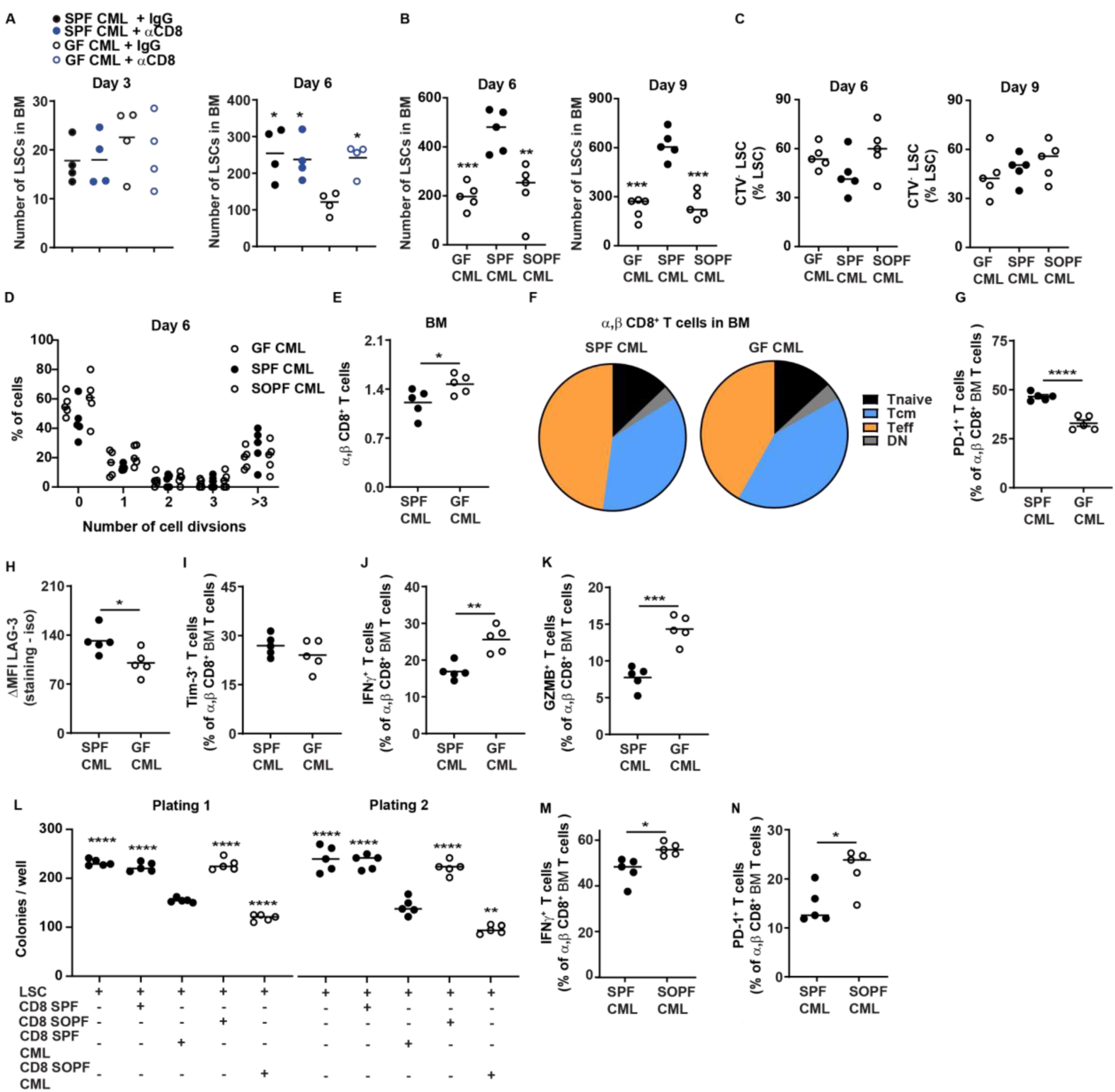
The gut microbiome inhibits CTL function in CML. (A-J) 3x10^4^ BL/6 LSCs were injected in GF and SPF mice or SOPF mice. (A) Numbers of LSCs in the BM of αCD8 antibody treated SPF and GF and control CML mice 3 and 6 days after CML induction (n=4 mice/group). Representative data from at least two independent experiments are shown. Significance was determined using a one-way ANOVA followed by Tukey’s post-test. (B) Numbers of LSCs and (C) frequency of proliferating LSCs and (D) cell divsions in the BM of SPF, GF and SOPF CML mice 6 and 9 days after CML induction (n=5 mice/group). Representative data from at least two independent experiments are shown. Significance was determined using a one-way ANOVA followed by Tukey’s post-test. mice. (E-K) 3x10^4^ LSCs were injected in GF and SPF mice. 15 days after injection, mice were sacrificed, and the BM CTLs were analyzed. (E) Frequencies of CTLs (n=5 mice/group). Significance was determined using a Mann Whitney test. (F) Differentiation state of CTLs (n=5 mice/group). Tnaive: CD8^+^CD44^-^ CD62L^+^; Tcm: CD8^+^CD44^+^CD62L^+^; Teff; CD8^+^CD44^+^CD62L^-^; DN: CD8^+^CD44^-^CD62L^-^. Significance was determined using a Mann Whitney test. (G) Frequency of PD-1-expressing CTLs. (H) ΔMFI of Lag-3 of CTLs. (I) Frequency of Tim-3-expressing CTLs. (J) Frequency of IFNℽ- and (K) granzyme B-producing CD8^+^α, β T cells. (n=5 mice/group). Significance was determined using a Mann Whitney test. (L, M) Colony and re-plating capacity of LSCs cultured in presence of CTLs derived from the BM of naive and CML SOPF and SPF mice (n=5 mice/group). Significance was determined using a one-way ANOVA followed by Dunnett’s multiple comparisons test (vs. CD8 SPF CML). (M, N) Frequency of (M) IFNℽ-producing and (N) PD-1 expressing CTLs in PP in BM of SPF and SOPF CML mice (n=5 mice/group). Representative data from at least two independent experiments are shown. Significance was determined using a Mann Whitney test. Results are shown as mean. *p < 0.05; **p < 0.01; ****p < 0.0001.

To determine how the gut microbiome affects CTLs in CML in the BM, we analyzed the number and function of CTLs in the BM of GF and SPF mice at a later stage of the disease (15 days after leukemia transplantation). We found that the BM of SPF CML mice contained significantly fewer CTLs compared to GF CML mice **(Fig. 5D)**_[OA1]_ In contrast, the differentiation state of the CTLs in BM were independent of the hygiene state of the mice **(Fig. 5E)**. Furthermore, CTLs from the BM of SPF mice expressed higher levels of the exhaustion markers PD-1 and Lag-3, but not Tim-3, on the cell surface **(Fig. 5F-H)**. The expression of these markers on CTLs in the BM of SPF mice was accompanied by a reduced capacity to produce the effector cytokine IFNℽ and granzyme B **(Fig. 5I, J)**.

We could previously demonstrate that BM CTLs in CML can kill LSCs *in vitro* via granzyme B^6^. To assess the potential of BM CTLs for their ability to eliminate LSCs at an early stage of the disease (day 8), we co-cultured LSCs with FACS-purified BM CTLs from naive and CML SOPF and SPF mice overnight followed by colony formation. BM CTLs from SPF CML mice were less potent in reducing LSCs *in vitro* compared to SOPF CML mice, as indicated by increased colony formation in serial re-plating experiments *in vitro* **(Fig. 5K)**. The lower capacity to eliminate LSCs was also reflected by reduced frequencies of activated IFNℽ and PD-1-expressing CTLs **(Fig. 5M, N)**.

Collectively, these data suggest that specific members of the gut microbiome can reduce CTL function in the BM of CML mice, leading to disease development.

## Discussion

Long-term eradication of leukemia can only be achieved by targeting LSCs that initiate and sustain the disease^2^. Despite TKI success in treating CML, dormant TKI-resistant LSCs remain in the BM in most patients and may lead to disease recurrence after drug withdrawal or through mutational acquisition^2^. For these patients, immunotherapy might be a potential therapeutic option. LSCs originate and expand in the BM close to naive and memory T cells^2,5^. CD4^+^ and CD8^+^ CTLs have been shown to contribute to immunosurveillance of leukemia, recognize leukemia associated antigens and lyse leukemia cells and LSCs^5^. However, LSCs also seem resistant to elimination by activated CD8^+^ CTLs *in vivo*, and various immune effector mechanisms have been reported to contribute to the expansion of LSCs rather than to their elimination^6,8,10,33,34^. The reasons why LSCs are selectively resistant to elimination by CD8^+^ CTLs remains poorly understood.

Gut bacteria have a highly variable interaction with the host, contributing to the regulation of immune cells in the gut and beyond during infection and cancer^12,35,36^. The degree of this interaction is regulated for example by the niche that bacteria colonize and whether the bacteria adhere to epithelial cells as well as the microbial products that they produce^37^. It is likely that certain microbial species, their antigens and secreted factors modulate the host immune system more than others. In the present study, we provide evidence that gut bacteria from the genus of *Sutterella* contribute to the development of CML by modulating the function of CD8^+^ CTLs in the BM. We found that colonization with the gram-negative, anaerobic, non-spore-forming bacteria from the genus *Sutterella*^26^ but not other gram-positive and gram-negative such *S. copri*, *L. reuteri* and *E.coli HS* is sufficient to promote CML development in SOPF mice by modulation of adaptive immune cells in the BM. Similarly, colonization with gram-negative *Bilophila*^27^ induced leukemia in SOPF mice. However, in contrast to all other bacterial strains tested in this study, colonization with *Bilophila* also increased the relative abundance of *Sutterella* spp. in SOPF mice. These data suggest *Sutterella* might be the main driver of leukemogenesis in the gut and that there might be synergy between *Bilophila* and *Sutterella* species in the gut. Furthermore, the effect of *Sutterella* could not be recapitulated in diverse gnotobiotic mouse models indicating that a complex and diverse microbiota is necessary to mediate the effect of *Sutterella* on leukemia development in our model.

*S. wadsworthensis* has been detected in 86% and 71% of adults and children, respectively^38,39^. *B. wadsworthia* was recovered in 60% of adult stool samples^40^. *Sutterella* spp. are abundant in the duodenum of healthy adults with a decreasing gradient toward the colon^31^. In addition, it has been frequently associated with human diseases, such as autism spectrum disorder, Down syndrome^27^, inflammatory bowel disease (IBD) and Crohn’s disease (CD)^41^. Recently, a study involving 10 patients with chronic lymphocytic leukemia (CLL) reported dissimilarities in stool microbiota abundance compared to healthy controls^42^. Of relevance for our study, they detected overrepresented *Bacteroides*, *Sutterella* and *Parabacteroides* in CLL relative to the average microbiota of healthy individuals. *Sutterella sp*. are considered to be mildly pro-inflammatory to the intestinal epithelium^31^, but do not drive inflammatory phenotype in IBD^39^. Others have reported *Sutterella sp*. as bystander species in intestinal disease due to their potential to degrade IgA^43^. However, our results show that *Sutterella sp.* also regulate the function of CTLs in the BM of CML mice, suggesting that *Sutterella sp.* have an immunomodulatory role in CML. Even though a role for the microbiome and specific gut bacteria in regulating anti-tumor immunity in CML has never been reported, there is ample evidence in solid tumor entities that the gut microbiota not only modulates anti-tumoral CTL immunity and that the composition of the intestinal microbiota is predictive for the efficacy of immune checkpoint therapy^15,16^. In addition, a recent study assessing the gut microbiota composition in 17 non-small cell lung cancer patients with an advanced disease who underwent more than three cycles of immune checkpoint therapy showed that *Bilophila* and *Sutterella* species were abundant in non-responders compared to responders^44^. Similarly, a comprehensive analysis of the microbiota of 775 patients with different types of solid cancers who received anti-PD-1, anti-PD-L1, or anti-CTLA-4 therapy associated the presence of the genera *Sutterella* and *Bilophila* with poor response to therapy^45^. Furthermore, a disturbed composition of the gut microbiota often results in a decreased abundance of *Firmicutes* and an increase of *Proteobacteria*, the phylum that also comprises *Sutterella* spp^46^.

Many molecules originating from the microbiota, such as bile acid metabolites or short-chain fatty acids have also been shown to shape the host immune system^35^. These factors are likely present in SPF mouse serum. However, repetitive SPF serum transfer could not restore CML development in SOPF mice in our model. In contrast, repeated administrations of *Sutterella* inactivated by peracetic acid but not heat-killed induced CML in SOPF mice, suggesting that sensing of *Sutterella* antigens by host immune/epithelial cells is necessary and sufficient for disease development in SOPF mice.

Antigen recognition in the gut is often mediated by the activation of Myd88/TRIF signaling on immune cells^37^. Similarly, Myd88/TRIF signaling has been shown to be important for the regulation of emergency hematopoiesis and to control self-renewal and differentiation of hematopoietic stem and progenitor cells under steady-state conditions, either directly or by modulating inflammatory cytokine production in the niche^18–21,47^. In our studies, we show that Myd88/TRIF signaling on host cells, but not on leukemia cells, promotes CML development in SPF mice. Furthermore, depletion of CTLs by antibody treatment rendered Myd88/Trif^-/-^ SPF mice susceptible to CML, again suggesting that antigen sensing is most likely to occur on CTLs or on professional antigen-presenting cells. This hypothesis is further supported by the finding that the BM of SOPF CML mice harbors a higher frequency of activated IFNℽ-producing CTLs with increased potential to kill LSCs *in vitro* compared to the BM of SPF CML mice. Josefdottir and colleagues have previously reported a role for the microbiota and T cells in regulating steady-state hematopoiesis^21^. Depletion of the gut microbiota by broad-spectrum antibiotic treatment altered T cell homeostasis in naive mice via disruption of Stat1 signaling resulting in impaired murine hematopoiesis. In addition, we found that CD8^+^ CTLs are more activated in the PP of SPF mice in contrast to the BM and identify *Sutterella* as a regulator of this phenotype. The difference in T cell activation between the two different organs may be explained by different levels of exposure to bacterial antigens and suggests that the microenvironment in which CTLs reside and the antigens to which they are exposed critically contribute to their activation state.

In conclusion, our study identifies bacteria from the genus *Sutterella* as central regulator of immune escape of LSCs and suggests that modulation of the gut microbiota may promote antileukemic immunity in CML.

## Acknowledgements

We thank the staff of the FACSlab (Department for BioMedical Research (DBMR), University of Bern, Switzerland) for providing excellent technical assistance and the staff of the central animal facility of the Medical Faculty, University of Bern, for their support. The Clean Mouse Facility is supported by the Genaxen Foundation, Inselspital and the University of Bern. This work was supported by grants from the Swiss National Science Foundation (310030_179394) and the Stiftung für klinisch-experimentelle Tumorforschung.

## Author contributions

MH performed experiments, analyzed and interpreted data, and contributed to the preparation of the Figures. FR designed and performed experiments, analyzed and interpreted data, and contributed to preparation of the figures and the writing of the manuscript. VR, CM, MR performed experiments, and analyzed and interpreted data. SCG, KDM, AJM and AFO interpreted data, designed experiments, and revised the manuscript. CR designed and supervised the study, interpreted data, and wrote the manuscript. All authors revised the manuscript and approved its final version.

## STAR Methods

Detailed methods include the following:

- Key Resources Table
- Resource Availability (Lead Contact, Materials availability, Data and code availability)
- Experimental models and subject details
- Methods Details
- Quantification and statistical analysis

## RESOURCES AVAILABILITY

### Lead contact

Further information and requests for resources and reagents should be directed to and will be fulfilled by the lead contact, Carsten Riether.

### Materials availability

All unique reagents generated in this study are available from the Lead Contact without restriction.

### Data and code availability

16S rRNA sequencing and metagenomics data will be made available at the time of publication under XXXX.

This study does not include the development of new code.

## EXPERIMENTAL MODEL AND SUBJECT DETAILS

### Mice

6 to 8 weeks old female C57BL/6J (BL/6) SPF and SOPF mice were purchased from Charles River Laboratories and were maintained on the same diet and in individually ventilated cages. C57BL/6J (BL/6), Germ-free, sDMDMm^24^ and Cuatro^25^ stably colonized mice were bred and maintained in flexible-film isolators at the Clean Mouse Facility, University of Bern, Switzerland as previously described^48^. Germ-free status was routinely monitored by culture-dependent and independent methods. All mice used in this study had access to food and water ad libitum and were regularly monitored for pathogens. Animal experiments were approved by the local experimental animal committee of the Canton of Bern (BE75/17, BE78/17, BE43/16, BE26/20, BE13/2021 and BE30/2021) and performed according to Swiss laws for animal protection.

## METHODS DETAILS

### Leukemia model

CML was induced and monitored as described before^6^. Briefly, FACS-purified LSKs from the BM of donor mice were transduced twice on two consecutive days with a BCR-ABL1-GFP retrovirus by spin infection. 3 x 10^4^ cells were injected intravenously into the tail vein of non-irradiated syngeneic recipients.

### CD8^+^α, β T cell depletion *in vivo*

Depletion of CD8^+^α, β T cells was performed by two injections of 75 µg αCD8 mAb (clone 2.43; BioXCell) at days -3 and -1 before leukemia induction.

### Colony-forming assays

For *in vitro* co-culture experiments, 10^3^ FACS-purified LSCs were incubated CD8^+^ T cells from BM of naive or CML SPF and SOPF mice at a ratio of 1:1 overnight in RPMI supplemented with 10% FCS (Thermo Fisher Scientific), 1% penicillin-streptomycin (MilliporeSigma), 1% L-glutamine (MilliporeSigma), SCF (100 ng/mL) and TPO (20 ng/mL) (Miltenyi Biotec), followed by plating in methylcellulose as previously described^6^. For each round of serial colony replating, total cells were collected from the methylcellulose, and 10^4^ cells were replated into methylcellulose without any T cells. Colony numbers were assessed with inverted light microscopy after 7 days for each round of plating (≥ 30 cells/colony).

### Bacterial culture

*Sutterella wadsworthensis* (DSM 14016) and *Bilophila wadsworthia* (DSM 11045) were cultured overnight in sterile anaerobic broth under static conditions (5 gr/l beef extract, 30 gr/l peptone, 5 gr/l yeast extract, 5 gr/l K2HPO, 40.5 gr/l L-Cysteine, 1 ug/ml hemin and 0.5 ug/ml VitaminK; with the addition of 1.6 g/l of sodium fumarate and 1.8 g/l of sodium formate for *S. wadsworthensis,* and taurine 0.025g/l for *B.wadsworthia). Segatella copri* (DSM 18205) was cultured overnight in sterile anaerobic broth under static conditions (TGM medium (Oxoid) with 5 gr/l beef extract, 1 ug/ml hemin and 0.5 ug/ml VitaminK). Anaerobic conditions were achieved in a Whitley MG500 anaerobic incubator gassed with 10% (v/v) H2, 10% CO2 and 80% N2. *L. reuteri* (DSM I49) was cultured in a closed glass bottle filled to about 80-90% vol with sterile MRS (Oxoid) broth and incubated overnight without shaking at 37°C. *Escherichia coli HS* was cultured overnight in sterile aerobic LB broth (5g NaCl, 5g yeast extract, 10g Trypton in 1 liter of distilled water) at 37 °C, shaking at 200 rpm. To prepare gavage solutions, bacteria were centrifuged for 10 min at 4,000*g* and washed twice with sterile PBS. The required dose (10^9^ CFU/ml) was resuspended in 300 μl of sterile PBS and administered to mice by oral gavage. Germ-free mice were gavaged upon one gavage, all the other mice under different hygiene conditions were colonised upon 4 consecutive gavages once every second or third day over 8-10 consecutive days.

### Preparation of heat-killed and peracetic acid-fixed bacteria

Overnight cultures of *S. wadsworthensis* and *B.wadsworthia* were collected, washed with PBS and resuspended at 10^9^ CFU/ml in PBS. For heat-killing, the aliquots were incubated for 10 minutes at 95°C on a pre-warmed heat block. For peracetic acid fixation^30^, peracetic acid (catalog: 433241, Sigma) was added to a final concentration of 0.6% in PBS. The suspension was mixed thoroughly and incubated for 1 h at RT. The bacteria were washed three times in 50 ml sterile D-PBS, carefully removing all supernatant after each centrifugation step, and thoroughly resuspending the pellet each time to remove the peracetic acid. The final pellet was resuspended at a final concentration of 10^9^ particles/ml in sterile D-PBS (determined by OD600).

300 μl of the killed bacteria stocks were administered to mice by oral gavage for 4 consecutive gavages once every second or third day over 8-10 consecutive days. The sterility of the killed bacteria was controlled by anaerobic and aerobic cultures.

### Serum transfer experiment

SPF C57BL/6 mice were terminally bled and 200 μl serum was injected i.v. undiluted in SOPF C57BL/6 recipients for 4 consecutive times every second or third day over 8-10 consecutive days.

### Fecal samples collection and DNA extraction

Fresh fecal pellets from mice were collected into sterile 2 mL Eppendorf tubes containing glass beads, for downstream fecal homogenization and microbial lysis. The collected fecal pellets were snap-frozen in liquid nitrogen, after which they were stored at −80°C until gDNA extraction. All fecal pellets within one experimental group were collected at the same time period. Fecal DNA was prepared using the QIAamp PowerFecal Pro DNA Kit (QIAGEN) according to the manufacturer’s instructions.

### 16S#rDNA IonTorrent sequencing

The microbiome analysis was performed according to the following protocol. Concentrations and purity of the isolated fecal DNA was evaluated by NanoDrop® (Thermo Scientific) and samples were stored at 4°C during library preparation and at −20°C thereafter for longer storage. The V5/V6 region of 16S rRNA genes was amplified with Platinum Taq DNA polymerase (Invitrogen) from 100 ng of fecal DNA using a range of oligonucleotide primers specific for the domains V5 and V6 of rDNA bacteria. Specifically, all forward core primers have been modified by the addition of a PGM sequencing adaptor, a ‘GT’ spacer and unique barcode that allow up to 96 different barcodes. The expected product length is 290 bp (350 bp including adaptors and barcodes). The different bacteria-specific primers were: IT_16S_FWD barcoded (5′-CCATCTCATCCCTGCGTGTCTCCGACTCAGC-barcode-ATTAGATACCCYGGTAGTCC-3′) and IT_16S_REV_1 (5′-CCTCTCTATGGGCAGTCGGTGATACGAGCTGACGACARCCATG-3′). Thermal cycling consisted of an initial 5 min at 94°C denaturation step, followed by 35 cycles of 1 min denaturation at 94°C, 20 s annealing at 46°C and 30 s extension at 72°C. Final extension consisted of 7 min at 72°C. PCR products after the first round of amplification were purified after 1% agarose gel electrophoresis by Gel Extraction Kit (QIAGEN). Purified amplicon concentration and purity was evaluated by quBit 3.0 Fluorometer (Thermo Fisher) prior to proceeding to the library preparation. The libraries were pooled at 26pM. To prepare template-positive Ion PGM™ Template OT2 400 Ion Sphere™ Particles (ISPs) containing clonally amplified DNA we used the Ion OneTouch™2 Instruments, with the Ion PGM™ Template OT2 400 Kit (for up to 400 base-read libraries) (Thermo Fisher). The template-positive ISPs was then enriched using the Ion OneTouch™ ES instruments (Thermo Fisher). In the end the sequencing was performed using the Ion PGMTM Sequencing 400 Kit with the Ion Personal Genome Machine (PGM™) System and the Ion 316™ Chip V2 (Thermo Fisher). All these instruments are part of the equipment of the Next Generation Sequencing platform of the University of Bern.

### Microbiome analysis

Using the QIIME pipeline, paired sequences were de-replicated and *de novo* as well as reference-based chimeras were removed using UCHIME. Sequences from all samples were merged, sorted by abundance, and closed operational taxonomic unit (OTU) picking at a threshold of 97% similarity was performed using USEARCH v5.2.236, followed by RDP classifier against the GreenGenes database for a stringent taxonomic assignment with a confidence interval of 80%. From the OTU abundance matrix and their respective taxonomic classifications, feature abundance matrices were calculated at different taxonomic levels (genus to phylum).

### Metagenomics analysis

Metagenomics analysis was performed at the PreBiomics S.r.l. 2024 (Italy) facilities.

#### Quality control

DNA samples were quantified by using the Quant-iT™ 1X dsDNA Assay Kits, BR (Life Technologies, #Q33267) in combination with the Varioskan LUX Microplate Reader (Thermo Fisher Scientific, #VL0000D0). Only samples that did exceed the threshold of 5 ng/μl were processed and diluted in RNase-DNase free water for the following library preparation.

#### Library preparation and sequencing

The sequencing libraries were prepared with the Illumina DNA Prep, (M) Tagmentation (96 Samples, IPB) kit (Illumina, #20060059) and the amplified libraries were purified with the double-sided bead purification procedure, as described by the Illumina protocol. Libraries concentration (ng/µl) was quantified with the Quant-iT™ 1X dsDNA Assay Kits, HS (Life Technologies, #Q33232) in combination with the Varioskan LUX Microplate Reader (Thermo Fisher Scientific, #VL0000D0). In addition, the base pair length (bp) was evaluated by using the D5000 ScreenTape Assay (Agilent, #5067-5588/9) in combination with the TapeStation 4150 (Agilent Technologies, #G2992AA). By knowing both library concentration and base pair length, it is possible to obtain the correct library volume to pool in the same tube in order to achieve optimal cluster density. The library pool was then quantified with the Qubit 1x dsDNA HS kit (Life Technologies, #Q33231) through the Qubit® 3.0 Fluorometer (Life Technologies, #Q33216) and the base pair length (bp) was evaluated as described before. Finally, the library pools were sequenced using the Novaseq Xplus platform (Illumina) with the NovaSeq X Series 25B Reagent Kit (300 Cycle) (Illumina, #20104706) at an average depth of 7.5Gbases per sample.

#### Preprocessing and quality control

Preprocessing and quality control of the sequenced samples were performed using a standalone pipeline available at https://github.com/SegataLab/preprocessing. Briefly, the software TrimGalore (version 0.6.6)^49^ was used for the read-level quality control step: reads with a quality score < 20, fragmented short reads (length < 75) and reads with more than 5 ambiguous nucleotides were removed. During the following screening for contaminant DNA, mouse DNA (GRCm39) and Illumina spike-ins PhiX DNA were removed using BowTie2 (version 2.3.4.3)^50^.

#### Taxonomic profiling

Taxonomic profiling was performed using MetaPhlAn (version 4.1.0) ^51^ against the Jun23 database with default parameters.

#### Statistical analysis

Alpha and beta diversity analyses were performed on the MetaPhlAn 4 profiles using the SciKit-bio (version 0.5.6), SciKit-learn (version 1.2.2) and SciPy (version 1.10.1) python libraries. Multidimensional Scaling (MDS) on arcsine square root-transformed abundances was performed using different distance metrics: Jaccard, Bray-Curtis and UniFra ^52^ PERMANOVA and Mann-Whitney tests were performed using the SciKit-bio python library.

#### Determination of the biomass

The weighed intestinal contents were suspended in 1 ml PBS at 30Hz shaking for 3 min. Fibre-filtered (100 μm) suspensions were diluted to OD700=∼0.8, stained with SYTO9 (5μM), and acquired at a Beckman Coulter MoFlo® ASTRIOS™ with known concentrations of spiked-in Fluoresbrite BB Carboxylate microsphere beads of various sizes (1 μm, 2 μm, Polyscience). Bacteria were identified as SYTO9+ and the bacterial identity of these populations was confirmed by using germ-free mice as negative controls. The bacterial concentration was calculated using the number of bacteria counted in the flow cytometer, the acquired volume as determined by the number of acquired beads, the dilution and the weight of the intestinal contents according to the equation below (mfec= weight of fecal pellet, Vres = volume to resuspend contents, cBac=concentration of bacteria per gram contents/feces, c1= concentration 1 μm beads, c2=concentration 2 μm beads, d1= dilution 1 μm beads, d2= dilution 2 μm beads, dbac=dilution bacteria, nbac=number of bacteria measured in flow cytometer, n1= number of 1 μm beads measured in flow cytometer, n2= number of 2 μm beads measured in flow cytometer)^53^.

#### FITC-dextran experiment

Before FITC-dextran administration, the mice were fasted for 4.5 hours in total (no food access, free water access). 25 mg/ml (500 mg/kg) in 200 microl/mouse of FITC-dextran 4 Kda (46944-100 SIGMA MG-F. Mol wt: 4000), diluted in PBS, was orally gavaged in some of the mice. After FITX-dextran administration mice were left without food and without water for 2.5 hours. Blood serum collection was conducted 2.5 hours after the FITC-dextran gavage. Blood serum was then diluted in 1:2 in PBS for FITC-measurment via Tecan plate reader (Excitation: 490 nm, Emission: 520 nm). Negative control mice were gavaged with water only, positive control mice were used after 5 days of Dextran-sodium sulfate 2% (DSS, 36-50 kDa; MP Biomedicals) administration in the drinking water.

#### Isolation of lymphocytes

To isolate lymphocytes from lymphoid tissues, spleens, lymph nodes and Peyer’s patches were cut into small pieces and digested in IMDM (2% fetal calf serum, FBS, Gibco) containing collagenase type IA (1 mg/ml, Sigma) and DNase I (10 U/ml, Roche) at 37°C for 30 min. Cellular suspensions were passed through a cell strainer (40 μm) and washed with IMDM (2% FCS, 2mM EDTA). Cells were centrifuged (600g, 7 min, 4°C) and resuspended in FACS buffer (PBS, 2 % FCS, 2mM EDTA, 0.01 % NaN_3_) for staining for flow cytometry analysis and counted at the Cytoflex instrument.

#### Flow cytometry

For surface staining, cells were washed once with DPBS before being stained with fixable viability dye (eBioscience) and FcR blocking reagent diluted in DPBS for 30 min on ice. Single cell suspensions were sequentially incubated with fluorescence-coupled antibodies diluted in FACS buffer for 15 min on ice.

For intracellular staining, cells were incubated at 37°C 5% CO2 with Phorbol 12-myristate 13-acetate (PMA, 250µg/ml, Invitrogen, catalog: J63916.MCR) and Ionomycin (250µg/ml, Sigma, catalog: I9657) in presence of Brefeldin A (2mg/ml, Sigma) for four hours. After that, cells were washed once with DPBS before being stained with fixable viability dye (eBioscience) and FcR blocking reagent diluted in DPBS for 30 min on ice. Single cell suspensions were sequentially incubated with fluorescence-coupled antibodies diluted in FACS buffer for 15 min on ice. Cells were then fixed and permeabilized using the Cytofix/Cytoperm Kit (BD Bioscience, catalog: 554714). Antibodies for intracellular staining were diluted in the permeabilization buffer from Cytofix/Cytoperm Kit and incubated at 4°C for 30 minutes.

The following mouse-specific conjugated antibodies were used: αCD8α-Brilliant Violet 785™ (clone 53-6.7, 1:1000, catalog 100749, RRID: AB_2562610), αCD44-APC/Cyanine7 (clone IM7, 1:400, catalog 103028, RRID: AB_830785), αCX3CR1-FITC (clone SA011F11, 1:50, catalog 149020, RRID: AB_2565703), αPD-1-PE (clone RMP1-30, 1:100, catalog 109104, RRID: AB_313421), αIFNg-Brilliant Violet 711™ (clone XMG1.2, 1:100, catalog 505835, RRID: AB_11219588), TNFα-PE/Dazzle™ 594 (clone MP6-XT22, 1:200, catalog 506346, RRID:AB_2565955), αLy6G/C-PE (clone RB6-8C5, 1:200, catalog 108408, RRID: AB_313373), αCD19-AlexaFluor700 (clone 6D5, 1:100, catalog 115528, RRID:AB_493735), αCD117-PE (c-kit, clone 2B8, 1:100, catalog 105807, RRID:AB_313216), αLy-6A/E-APC (Sca-1, clone D7; 1:100, catalog 108111, RRID:AB_313349), αCD117-APC-Cy7 (c-kit, clone 2B8, 1:300, catalog 105838, RRID:AB_2616739), αCD19-APC-Cy7 (clone 6D5, 1:300, catalog 115530, RRID:AB_830707), αCD4-BV650 (clone RM4-5, 1:600, catalog 100555, RRID:AB_2562529), αCD8–Alexa Fluor 700 (clone 53-6.7, 1:800, catalog 100729, RRID:AB_493702), αCD62L–Pacific Blue (clone MEL-14; 1:200, catalog 104423, RRID:AB_493381), αCD366-APC (Tim-3, clone: RMT3-23, 1:100, catalog 119706, RRID:AB_2561656), αCD279-Brilliant Violet 421 (PD-1, clone: RPM1-30, 1:200, catalog 109121, RRID:AB_2687080) were purchased from BioLegend. viability dye e450 (1:4000) were purchased from Thermo Fisher Scientific.

Lin^+^ cells were excluded by magnetic-activated cell sorting (MACS) using biotinylated αCD19 (clone 6D5, 1:300, catalog 115504, RRID:AB_313639), αCD3e (clone 145-2C11, 1:300, catalog 100304, RRID:AB_312669), αLy-6G/C (clone RB6-8C5, 1:300, catalog 108404, RRID:AB_313369), and αTer119 (clone Ter-119; 1:300, catalog 116203, RRID:AB_313704) from BioLegend, followed by a second staining step with streptavidin Horizon-V500 (1:1000, catalog 561419, RRID:AB_10611863) from BD Biosciences after the separation.

Data were acquired on a LSR-Fortessa (BD Biosciences) and analyzed using FlowJo software v10 (Tree Star Inc.). In all experiments, FSC-H versus FSC-A was used to gate singlets, dead cells were excluded using the fluorescence-coupled fixable viability dye (eBioscience).

#### Statistical analysis

All flow cytometry, in vitro and in vivo data were analyzed and plotted using GraphPad Prism® software v9.0 (GraphPad). Bars and error bars indicate means and standard deviations of the indicated number of independent biological replicates. Two-tailed Student’s t test, Mann-Whitney test, one-way-ANOVA followed by Tukey’s post-test and two-way ANOVA followed by Bonferroni’s post-test were used as indicated in the figures legends. Significant differences in Kaplan-Meier survival curves were determined using the log-rank test. Data are represented as mean. P<0.05 was considered significant. Details on the quantification, normalization and statistical tests used in every experiment can be found in the corresponding figure legend. n represents the number of independent replicates in each experiment.

