## Supplemental Figures for "Low abundant intestinal commensals modulate immune control of chronic myeloid leukemia stem cells"

**Supplemental Figures and legends**

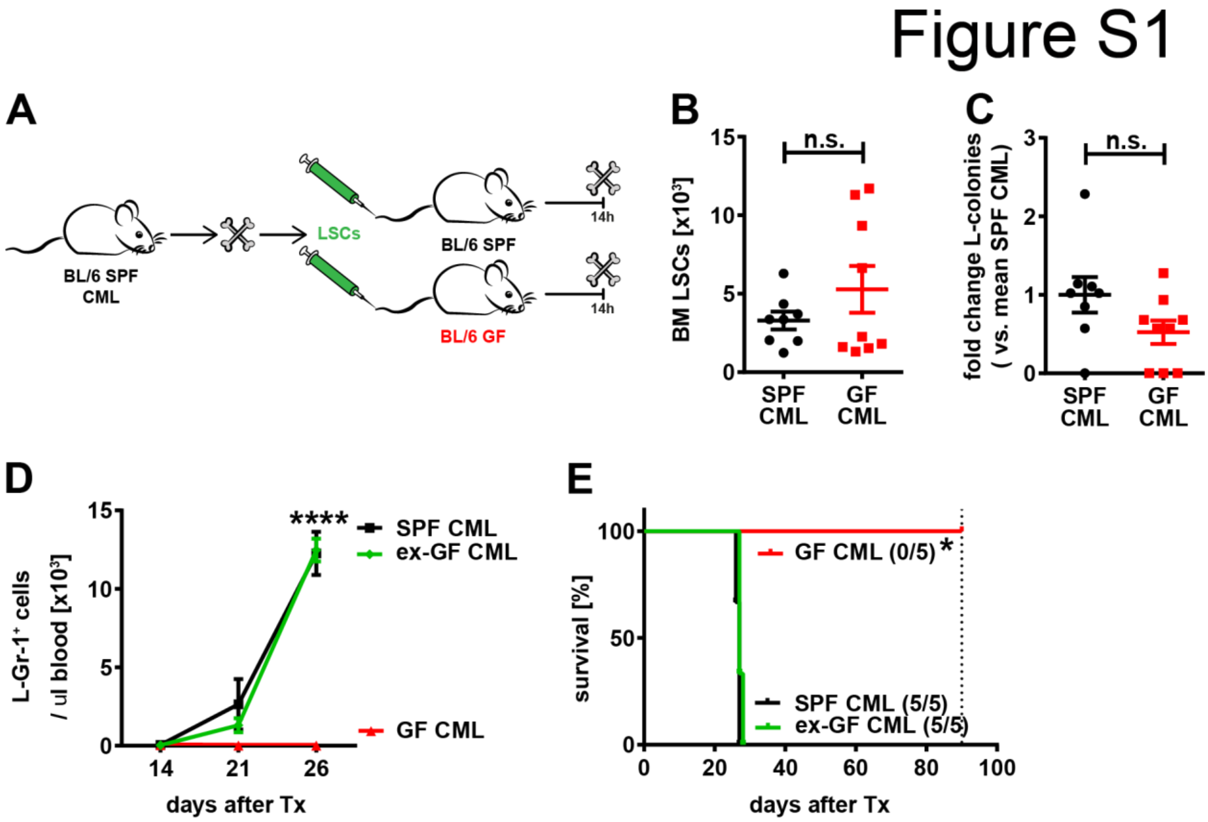


**Figure S1: Colonization with SPF microbiota restores CML development in GF mice.** (A) Experimental plan. 3x$10^4$ LSCs were injected i.v. into non-irradiated C57BL/6 SPF (SPF) and germ-free (GF) recipient mice and mice were sacrificed 14h later. (B) Numbers of LSCs and (C) Fold-change in BCR-ABL1-GFP$^+$ colony formation in BM plated SPF mice and GF mice previously injected with LSCs (n=8-9 mice/group). Pooled data from at least two independent experiments are shown. Significance was determined using a Student's t-test. (D) Numbers of BCR-ABL1-GFP$^+$ granulocytes/μl in blood (E) and Kaplan-Meier survival curves resulting from primary transplantations (Tx) of LSCs in untreated C57BL/6 SPF and GF mice and GF mice after co-housing for 4 weeks with SPF mice (ex-GF mice; n=5 mice/group). Representative data from at least two independent experiments are shown. Significance was determined using a log-rank test. Data are represented as mean ± SD. *$p < 0.05$; ****$p < 0.0001$.

### Figure S2

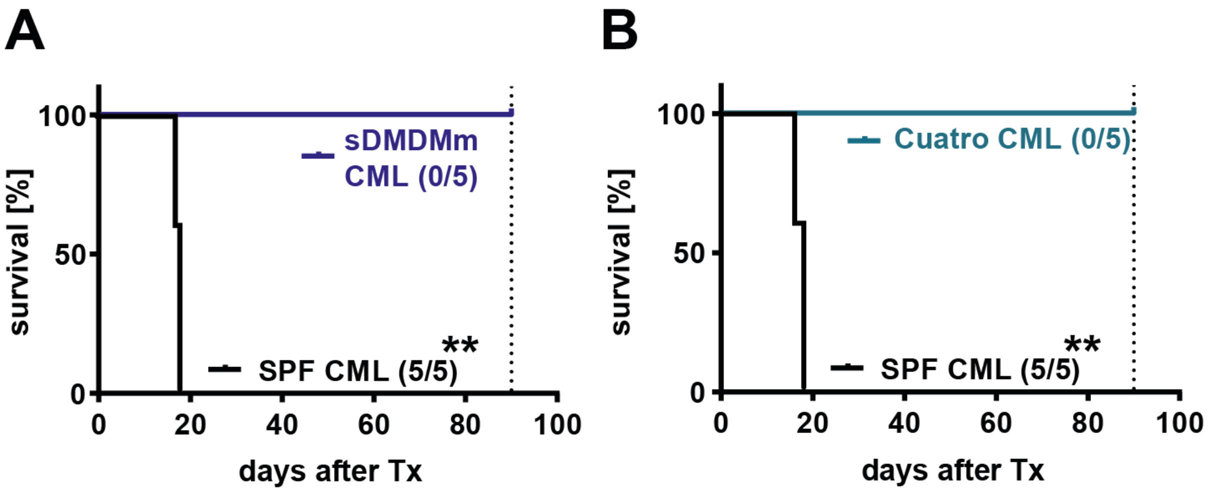


**Figure S2: Gnotobiotic sDMDMm- and Cuatro-colonized mice are protected from CML development.** Kaplan-Meier survival curves resulting from primary transplantations (Tx) of $3x10^4$ LSCs in C57BL/6 SPF and (A) sDMDMm or (B) Cuatro mice (n=5 mice/group). Representative data from at least two independent experiments are shown. Significance was determined using a log-rank test. **p < 0.01.

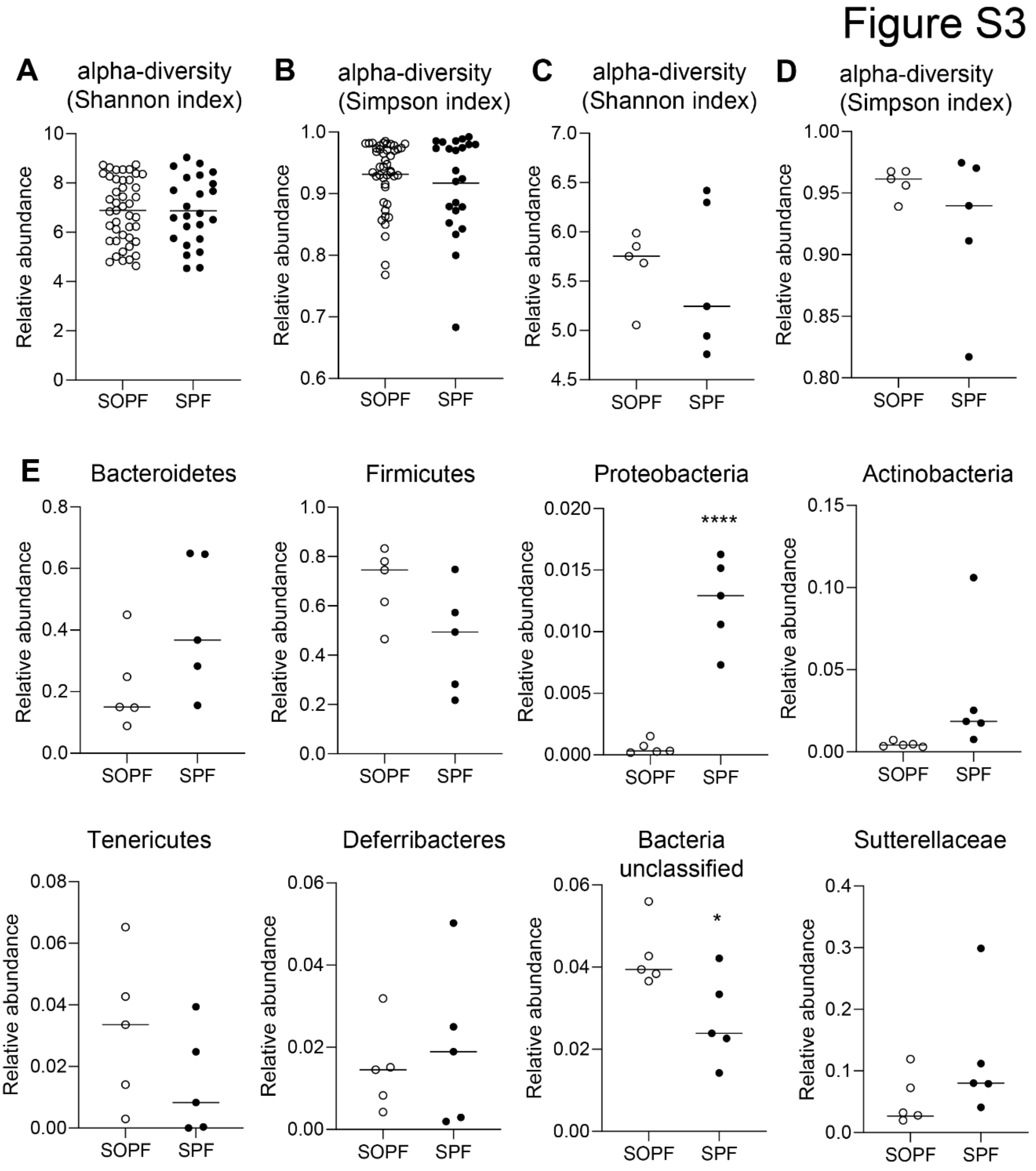


**Figure S3: Species diversity from ionTorrent analysis and metagenomics analysis of fecal samples from SOPF and SPF mice**. (A-B) Microbiota composition analysis with 16S rRNA amplicon sequenicng with ionTorrent. Species diversity of fecal samples from SPF and SOPF mice (n=44 and 23, respectively). (C-E) Metagenomics analysis of fecal samples from SOPF and SPF mice (n=5/group). (C-D) Species diversity: (C) Shannon index. (D) Simpson index. (E) Relative abundances of indicated bacteria taxa in feces from C57BL/6 SOPF and SPF mice. (n=5/group). Significance was determined using a Mann Whitney test. Results are shown as mean. *p < 0.05; ****p < 0.0001.

#### Figure S4

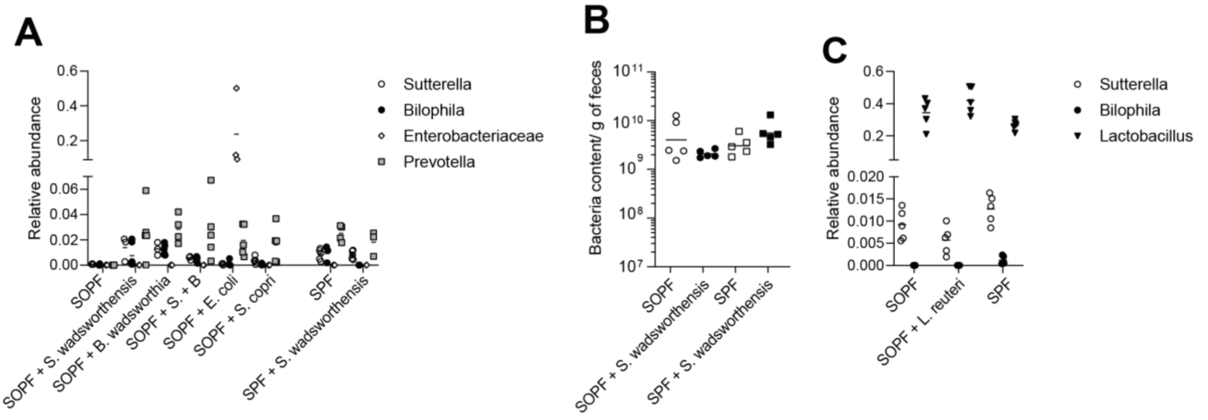


**Figure S4: Relative bacteria abundance and biomass.** (A) Relative abundance of indicated bacteria taxa in feces from naive SPF and SOPF mice or mice after gavage with *S. wadsworthensis*, *B. wadsworthia*, *S. copri* and *E. coli* (n= 3-6 mice/group). (B) Bacterial amount per gram of feces (biomass) in SPF and SOPF naive and after gavage with S. *wadsworthia* (n=5 mice/group). (C) Relative abundance of indicated bacteria taxa in feces from naive SPF and SOPF mice or SOPF mice after gavage with *L. reuteri* (n=5 mice/group).

#### Figure S5

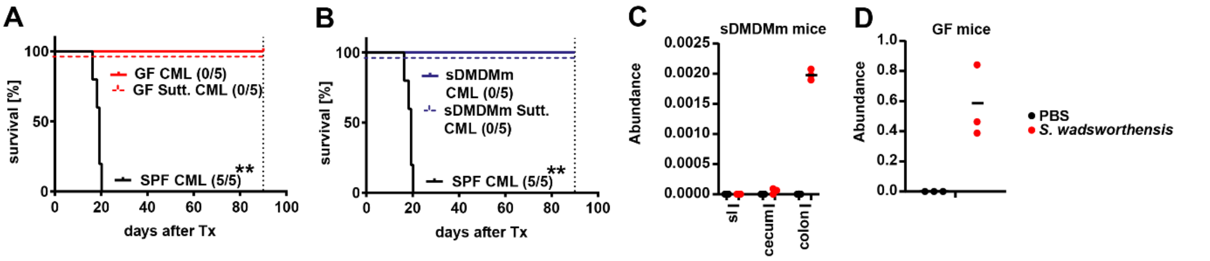


**Figure S5: Colonization with S. *wadsworthensis* does not promote CML development in GF and sDMDMm mice.** (A-D) C57BL/6 GF or sDMDMm mice were colonized with *S. wadsworthensis* at days -10, -7, -3, and 0 prior to CML induction. Kaplan-Meier survival curves resulting from primary transplantations (Tx) of $3x10^4$ LSCs in C57BL/6 SPF and (A) GF mice or (B) sDMDMm (n=5 mice/group). (C) Relative abundance of *S. wadsworthensis* in the small intestine (sI), cecum and colon of sDMDM mice or (D) in feces of GF mice (n=3 mice/group). Representative data from at least two independent experiments are shown. Significance was determined using a log-rank test. **p < 0.01.

### Figure S6

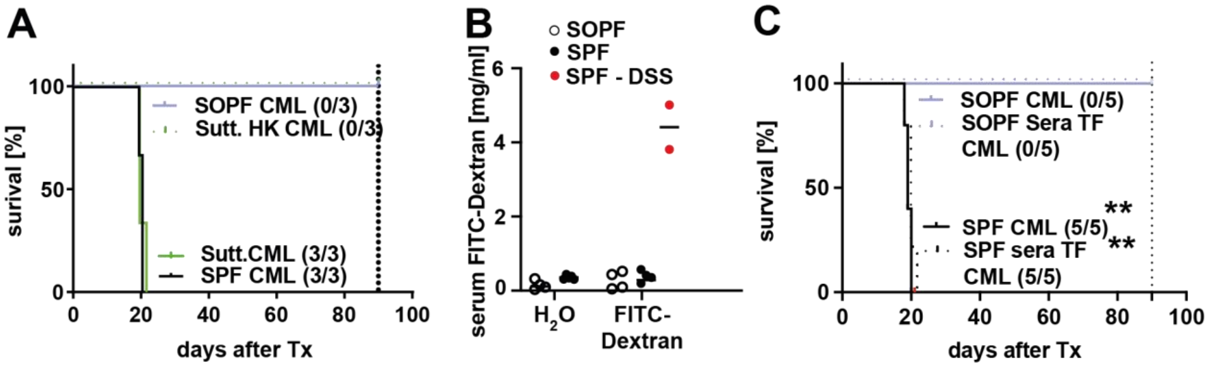


**Figure S6: Presence of inactivated intact bacterial cells , but not circulating bacterial products or metabolites, promote development of CML in SPF mice.** (A) Kaplan-Meier survival curves of SPF and SOPF CML mice colonized with live and heat-killed bacteria (HK) *S. wadsworthensis* (n=3 mice/group). Representative data from at least two independent experiments are shown. (B) FITC-dextran concentrations in peripheral blood of SOPF, SPF and SPF-DSS mice 2.5 hours after oral application (n=2-4 mice/group). (C) Kaplan-Meier survival curves of SPF and SOPF CML mice that were repetitively injected with vehicle or sera from SPF mice prior to leukemia induction (Tx) (n=5 mice/group). Representative data from at least two independent experiments are shown. Significance was determined using a log-rank test. Data are represented as mean. $^{**}p < 0.01$.

### Figure S7

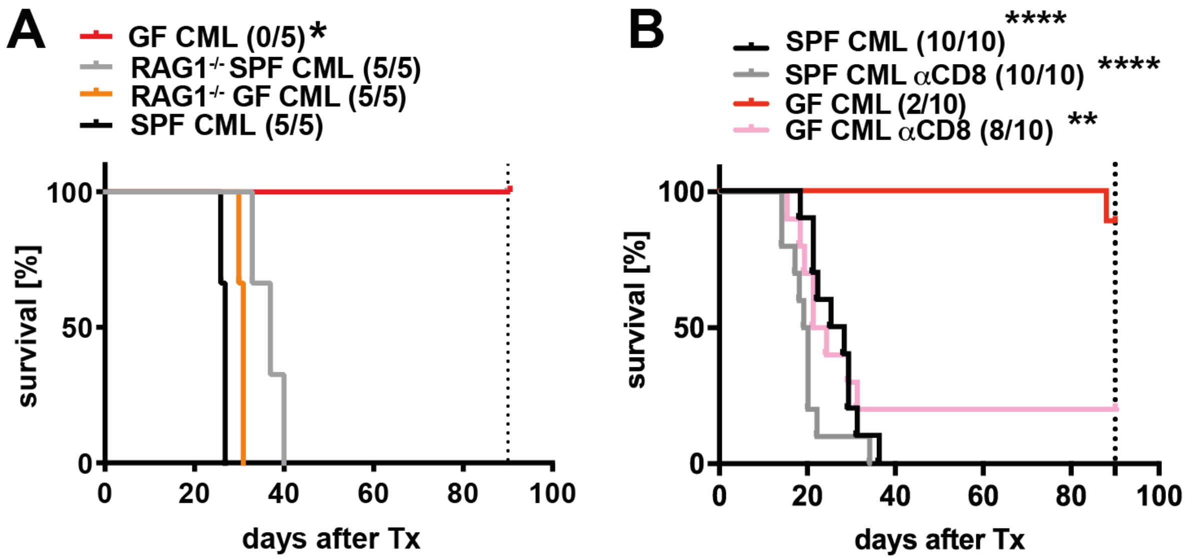


**Figure S7: CML develops in GF mice lacking CD4⁺ and CD8⁺ and in GF lacking CTLs.** Kaplan-Meier survival curves resulting from primary transplantations of LSCs into RAG-1$^{-/-}$ mice (A, n=5 mice/group) and αCD8 antibody-treated SOPF mice (B, n=10 mice/group). Representative data from at least two independent experiments are shown. Significance was determined using a log-rank test. *$p < 0.05$; **$p < 0.01$; ****$p < 0.0001$.
